# REBASTINIB, OLVEREMBATINIB, AND NAS-181 INHIBIT *MYCOBACTERIUM ABSCESSUS* BY DISRUPTING MYCOTHIOL HOMEOSTASIS

**DOI:** 10.1101/2025.08.22.671851

**Authors:** David L. Serrano, Elena M. Duarte, Arjun Mehta, Robert J. Collins, A. P. Sugunan

## Abstract

Nontuberculous mycobacteria (NTM), particularly *Mycobacterium abscessus*, pose a major clinical challenge due to intrinsic drug resistance and limited therapeutic options. In this study, we evaluated three repurposed kinase inhibitors, such as rebastinib, olverembatinib, and NAS-181for their antimycobacterial potential against *M. abscessus*. All three compounds displayed measurable in vitro activity, with MIC values of 22 μM, 40 μM, and 9 μM, respectively. Whole-genome sequencing of resistant mutants revealed nonsynonymous mutations in key enzymes of the mycothiol biosynthesis pathway, including D87A and W227G substitutions in *MshC* (*MAB_2116*) and a Q51T substitution in *MshA* (*MAB_4057c*), implicating disruption of redox homeostasis as a likely mechanism of action. Consistent with this, biochemical analyses demonstrated a concentration-dependent decrease in intracellular mycothiol (MSH) and its precursor GlcN-Ins upon treatment with each compound. Cytotoxicity profiling indicated tolerable activity thresholds (rebastinib, 30 μM; olverembatinib, 60 μM; NAS-181, 14 μM), enabling further macrophage infection assays. All three compounds significantly reduced intracellular *M. abscessus* burden in THP-1 macrophages without affecting host cell viability. In a mouse pulmonary infection model, treatment with rebastinib, olverembatinib, or NAS-181 (5 mg/kg) resulted in maintenance of body weight, decreased lung CFU counts, and reduced spleen enlargement compared to vehicle controls, with efficacy comparable to amikacin. Collectively, these findings identify rebastinib, olverembatinib, and NAS-181 as promising repurposed drug candidates that target mycothiol biosynthesis, providing a foundation for the development of novel therapeutic strategies against drug-resistant *M. abscessus*.

## 1. INTRODUCTION

Infections caused by nontuberculous mycobacteria (NTM) are increasingly acknowledged as a serious and growing health concern ^1^. Among the rapid-growing NTM species, *M. abscessus* is the most frequently isolated pathogen in clinical settings ^2–4^. It primarily causes pulmonary disease in individuals with underlying immune deficiencies or preexisting lung disorders, such as cystic fibrosis or bronchiectasis ^2^. Although it was long thought that *M. abscessus* infections were solely acquired from environmental sources like soil and water, recent evidence has revealed person-to-person transmission, particularly among cystic fibrosis patients ^1,5,6^.

Taxonomically, *M. abscessus* is complex and has undergone several reclassifications. Large-scale genomic studies now clearly divide the species into three subspecies—*M. abscessus* subsp. *abscessus*, subsp. *bolletii*, and subsp. *massiliense*—superseding earlier two-subspecies classifications ^7^.

Treatment of *M. abscessus* infection remains extremely challenging due to the organism’s high intrinsic resistance to most antibiotics, including those used against *M. tuberculosis* ^8,9^. Clinical guidelines recommend maintaining negative sputum cultures for at least 12 months during therapy, which means patients often undergo treatment with at least three drugs for several years ^1,10^. Because chemotherapy alone performs poorly, surgical removal of infected lung tissue is advised when feasible ^1^. Standard treatment regimens include macrolides (typically clarithromycin) together with parenteral agents such as amikacin, imipenem, or cefoxitin ^1^. Therapeutic success is further hampered by the widespread emergence of inducible clarithromycin resistance, which is largely mediated by the erm(41) gene. Although this gene is present in all subspecies, inducible macrolide resistance mainly occurs in subsp. *abscessus* and subsp. *bolletii*, while subsp. *massiliense* contains nonfunctional erm(41) alleles ^11^. Notably, not all subsp. *abscessus* strains express inducible resistance; some lack it due to specific polymorphisms in erm(41) ^12^.

Despite the clear medical need, progress in developing new drugs for NTM infections has been slow ^13^. One potential approach is repurposing existing drugs with unrecognized anti-*M. abscessus* activity^14^. Drug repurposing has proven effective for other difficult-to-treat bacterial pathogens and could provide a faster route toward new therapies. A previous study, for example, suggested that metronidazole exhibited potent activity against *M. abscessus*, though this finding could not be independently confirmed ^15^.

In our small-molecule screening campaign based on the Clinical Compound Library Plus (MCE), a curated collection of approximately 3,468 clinical-stage compounds designed for high-throughput (HTS) and high-content screening (HCS), we identified several compounds exhibiting antimycobacterial activity. Three of hits were rebastinib, olverembatinib, and NAS-181. These compounds were originally developed as tyrosine kinase inhibitors for cancer therapy ^16–19^, but in vitro assays performed during our screening revealed significant growth-inhibitory effects against clinically relevant mycobacterial strains, including drug-resistant isolates. This unexpected finding suggests that rebastinib, olverembatinib, and NAS-181may be repurposed as antimycobacterial agents and provides a rationale for further optimization of these scaffolds to improve potency and selectivity.

## 2. METHOD

### 2.1. Compounds

A total of 3,468 compounds from the Clinical Compound Library Plus (MedChemExpress) were screened in this study. This library consists of clinical-stage molecules and FDA-approved drugs, which makes it suitable for drug repurposing campaigns. Compounds were supplied as pre-solubilized stocks in DMSO and dispensed directly into assay plates at the indicated screening concentration. For confirmation of primary hits identified in the initial screen, rebastinib, olverembatinib, and NAS-181 were re-purchased individually from commercial sources (MedChemExpress) and freshly prepared in DMSO according to the manufacturer’s instructions. All other reference drugs used in this work were purchased from Sigma.

### 2.2. Bacteria

The study was conducted with *M. abscessus* strain ATCC 19977. Liquid cultures were grown in Middlebrook 7H9 broth (BD Difco) supplemented with 0.5% albumin, 0.2% glucose, 0.085% sodium chloride, 0.0003% catalase, 0.2% glycerol, and 0.05% Tween-80. Solid media consisted of Middlebrook 7H10 agar (BD Difco) supplemented with 0.5% albumin, 0.2% glucose, 0.085% sodium chloride, 0.5% glycerol, 0.0003% catalase, and 0.006% oleic acid.

### 2.3. Single-point growth inhibition screening assay

The Clinical Compound Library Plus was screened using a 96-well microdilution assay. Briefly, compounds were tested at a single-point concentration of 20 µM in flat-bottom 96-well plates (Corning Costar) with a starting inoculum of *M. abscessus* ATCC 19977 at OD_600_ = 0.05 (∼10^7^ CFU/ml) in a final volume of 200 µl per well. The inoculum was prepared from a mid-log phase preculture (OD_600_ 0.4–0.6). Plates were sealed with a Breathe-Easy™ membrane (Sigma-Aldrich), placed in an airtight humidified container, and incubated for 3 days at 37 °C on an orbital shaker (110 rpm). Each plate included a medium-only control, a drug-free growth control, and a positive control (clarithromycin, 20 µM). Following incubation, the cultures were manually resuspended, and OD_600_ values were measured using a TECAN Infinite Pro 200 plate reader. Compounds that produced ≥80% growth inhibition relative to the untreated control were defined as hits. All experiments were performed in triplicate.

### 2.4. MICs determination

Minimum inhibitory concentrations (MICs) of the selected compounds were determined using the resazurin microtiter assay. Briefly, exponential-phase *M. abscessus* cultures were adjusted to an OD_600_ of 0.01, and 50 µL of bacterial suspension were dispensed into each well of a sterile 96-well microtiter plate. Serial 2-fold dilutions of the test compounds (50 µL) were added to the wells, resulting in a total volume of 100 µL per well. Drug-free growth controls were included on each plate. To minimize evaporation, 200 µL of sterile water was added to all edge wells. Plates were sealed with self-adhesive membranes and incubated at 37 °C for 3 days. Subsequently, 40 µL of 0.025% (w/v) resazurin solution were added to each well and plates were further incubated overnight. Fluorescence was measured using a TECAN Infinite Pro 200 plate reader. Dose– response curves were generated and MIC_50_ values were calculated using GraphPad Prism (GraphPad Software, San Diego, CA, USA).

### 2.5. Resistant mutant frequency determination

To determine the frequency of spontaneous resistance, 250 µL of exponential-phase *M. abscessus* were plated onto Middlebrook 7H10 agar containing 0, 1×, 2×, 4×, or 8× MIC_90_ of the respective test compounds. Plates were incubated for 5 days at 37 °C, and the number of resistant colonies was recorded. The mutation frequency was calculated as the median number of resistant colonies divided by the total number of viable bacteria plated.

### 2.6. Checkerboard interaction assay

Compound interactions were assessed using a resazurin-based checkerboard assay. Two-fold serial dilutions of rebastinib, olverembatinib, and NAS-181 were combined with standard anti-*M. abscessus* drugs in 96-well microplates, starting at 8× MIC_50_ for each compound. Exponential-phase *M. abscessus* CIP104536T cultures (OD_600_ = 0.0025) were added to each well and plates were incubated for 5 days at 30 °C. Resazurin was then added, and fluorescence was measured after overnight incubation. Fractional inhibitory concentration indices (ΣFIC) were calculated to evaluate the interactions: ≤0.5 = synergy, >0.5–4 = additive, and >4 = antagonistic. To confirm the results, aliquots from each well were plated on 7H10 agar and CFU were counted after incubation at 37 °C for 3 days.

### 2.7. Identification of genetic variants

Genomic DNA was isolated from the parental *M. abscessus* ATCC 19977 strain and resistant mutants generated in the presence of rebastinib, olverembatinib, and NAS-181 using bead beating followed by phenol:chloroform:isoamyl alcohol extraction. Whole-genome sequencing of the parental strain and the selected resistant isolates was performed using Illumina HiSeq 2500 technology. Raw fastq files were trimmed with PRINSEQ-lite (version 0.20.4) using the following parameters: -min_len 50, -min_qual_mean 30, -trim_qual_right 30, -ns_max_n 0, and -noniupac. Quality-filtered reads were aligned to the *M. abscessus* ATCC 19977 reference genome using BWA-MEM. Alignment files were converted to BAM format and sorted using SAMtools, and duplicate reads were removed with Picard MarkDuplicates. GATK HaplotypeCaller was used to identify SNPs and indels, and resulting variants were annotated with snpEff. Variants with a sequencing depth ≤5 or located in repetitive PE/PPE/PE_PGRS regions were excluded from the analysis. Selected mutations were confirmed by Sanger sequencing.

### 2.8. Cytotoxicity assay

THP-1 cells were differentiated with phorbol 12-myristate 13-acetate (PMA) for 48 h and subsequently infected with *M. abscessus* at a multiplicity of infection (MOI) of 1 for 3 h at 37 °C in 5% CO_2_. After infection, extracellular bacteria were eliminated by treatment with 250 µg/mL amikacin for 2 h. Cells were washed with PBS and exposed to serially decreasing concentrations of test compounds for 72 h at 37 °C in 5% CO_2_. Following incubation, cells were lysed and serial tenfold dilutions of the lysates were plated onto Middlebrook 7H10-OADC agar. Plates were incubated at 37 °C for 3–4 days, and CFU were counted to determine intracellular bacterial survival.

### 2.9. Intracellular bacterial replication assay

To evaluate the intracellular activity of the compounds, THP-1 macrophages were seeded in 96-well plates and infected with *M. abscessus* at a multiplicity of infection (MOI) of 1 for 3 h. After infection, extracellular bacteria were removed by washing and killed by treatment with amikacin (250 µg/mL for 2 h). Cells were washed with PBS and subsequently treated with serial dilutions of the test compounds for 3 days. Infected cells were then lysed, and the lysates were serially diluted in PBS and plated onto Middlebrook 7H10-OADC agar. Plates were incubated at 37 °C for 3–4 days, and colony-forming units (CFU) were enumerated to assess intracellular bacterial survival.

### 2.10. Determination of mycothiol content

*M. abscessus* was cultured in Middlebrook 7H9 medium in the absence of compound (control) or in the presence of 1×, 2×, 4×, or 8× MIC_90_ of rebastinib, olverembatinib, or NAS-181. Cultures were harvested at exponential phase and bacterial cells were collected by centrifugation. Cell pellets were resuspended in 50% acetonitrile / 20 mM HEPES (pH 8.0), with or without 2 mM monobromobimane (mBBr), and incubated at 60 °C for 15 min to extract cellular thiols. Debris was removed by centrifugation and the supernatants were stored at –70 °C until analysis. Mycothiol (MSH) was quantified by derivatization with mBBr to form the mycothiol–bimane (MSmB) adduct, followed by fluorescence HPLC analysis. Results were expressed as nmol per 10^9^ cells using the conversion 1 OD_600_ = 2.5 × 10^8^ cells.

### 2.11. Animal study

Seven-week-old female BALB/c mice (∼20 g) were housed under specific pathogen-free (SPF) conditions with a 12 h light/12 h dark cycle, a temperature of 18–23 °C, and 40–60% relative humidity. Mice were acclimatized for 7 days prior to the start of the experiment.

A pulmonary *M. abscessus* infection model was established by intraperitoneal administration of cyclophosphamide (150 mg/kg) on day −4 and day −1 (in 0.2 mL PBS) to induce transient immunosuppression. On day 0, mice were anesthetized with 2–5% isoflurane and infected intranasally with 1 × 10^6^ CFU of a clinical *M. abscessus* rough morphotype isolate (ID No. 5) in 40 μL PBS.

Starting 24 h post infection, mice were randomly divided into five treatment groups (n = 5 per group) and received daily intraperitoneal injections of either vehicle (PBS, 0.1 mL), 5 mg/kg rebastinib, 5 mg/kg olverembatinib, 5 mg/kg NAS-181, or 50 mg/kg amikacin. All treatments were administered for 12 consecutive days.

On day 12, mice were euthanized under anesthesia by cervical dislocation. Spleens were excised, blotted dry, and weighed, and the spleen weight was normalized to the corresponding body weight. The postcaval lobe of the lung was collected and homogenized in PBS. Lung homogenates were serially diluted and plated on Middlebrook 7H10 agar, incubated at 37 °C, and colony-forming units (CFU) were enumerated to determine the lung bacterial burden.

### 2.12. Statistical analyses

Statistical analyses were performed on Prism 5.0 (GraphPad) and detailed for each figure legend. *, P ≤ 0.05, **, P ≤ 0.01, ***, P ≤ 0.001, ****, P ≤ 0.0001.

## 3. RESULT

### 3.1. Screening of clinical compounds identifies rebastinib, olverembatinib, and NAS-181 as inhibitors of *M. abscessus*

To identify potential compounds with antimycobacterial activity, we screened the Clinical Compound Library Plus (3,468 compounds) at 40 µM against the reference strain *M. abscessus* ATCC 19977. Using a cutoff of ≥80% growth inhibition, we identified three primary hits (0.19% hit rate). Three of them included the tyrosine kinase inhibitors rebastinib (RES), olverembatinib (OLV), and NAS-181 and further studied in the following expreiment. The primary hits were subsequently repurchased and tested in dose–response assays, which confirmed their antimycobacterial activity. Rebastinib and olverembatinib are small-molecule tyrosine kinase inhibitors originally developed for cancer therapy, while NAS-181 is a selective 5-HT_2_A receptor antagonist. None of these compounds were previously reported to possess antimycobacterial activity, highlighting their potential for drug repurposing against *M. abscessus*.

### 3.2. Rebastinib, olverembatinib, and NAS-181 are active against reference strains representing all three subspecies of the *M. abscessus* complex

To evaluate whether the identified hits display consistent activity across the phylogenetic spectrum of the *M. abscessus* complex, we determined their MIC_50_ values against the reference strains *M. abscessus* subsp. *abscessus, M. abscessus* subsp. *bolletii*, and *M. abscessus* subsp. *massiliense*. All three compounds showed measurable inhibitory activity (Table 1). Rebastinib exhibited MIC_50_ values ranging from 17 to 30 µM, olverembatinib from 35 to 42 µM, and NAS-181 from 7 to 12 µM. These results indicate that all three compounds retain activity across the *M. abscessus* subspecies, with NAS-181 showing the most potent inhibitory effect.

**Table 1.**
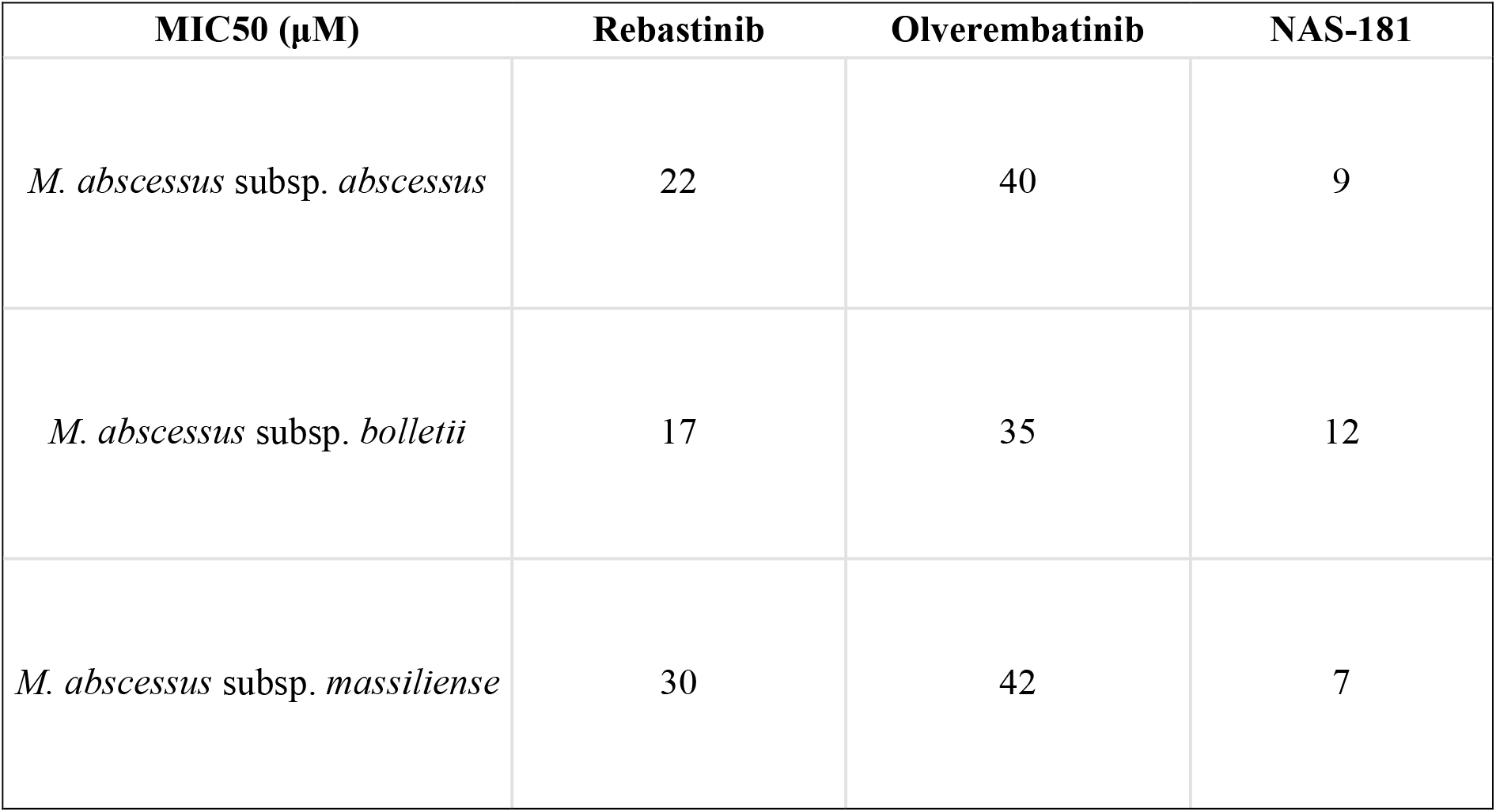
MIC_50_ values of rebastinib, olverembatinib, and NAS-181 against reference strains of the *M. abscessus* complex.

### 3.3. Resistance frequency comparison between rebastinib, olverembatinib, NAS-181, and clarithromycin

To compare the resistance development potential of the three candidate compounds with clarithromycin (CLA), we measured resistant mutant frequencies at different multiples of the MIC (1×, 2×, 4×, and 8× MIC) on Middlebrook 7H10-OADC agar. The frequencies of resistant mutants for rebastinib ranged from 8.0 × 10^−6^ to 3.0 × 10^−7^, and for olverembatinib from 7.2 × 10^−6^ to 2.2 × 10^−8^ (Table 2). By contrast, NAS-181 displayed a markedly lower resistance frequency, ranging from 2.5 × 10^−7^ to 3.5 × 10^−8^, which was closer to the values observed for clarithromycin (6.0 × 10^−8^ to 6.0 × 10^−9^). These findings indicate that while rebastinib and olverembatinib are associated with higher frequencies of spontaneous resistance, NAS-181 demonstrates a lower resistance liability, comparable to that of the standard drug clarithromycin.

**Table 2.**
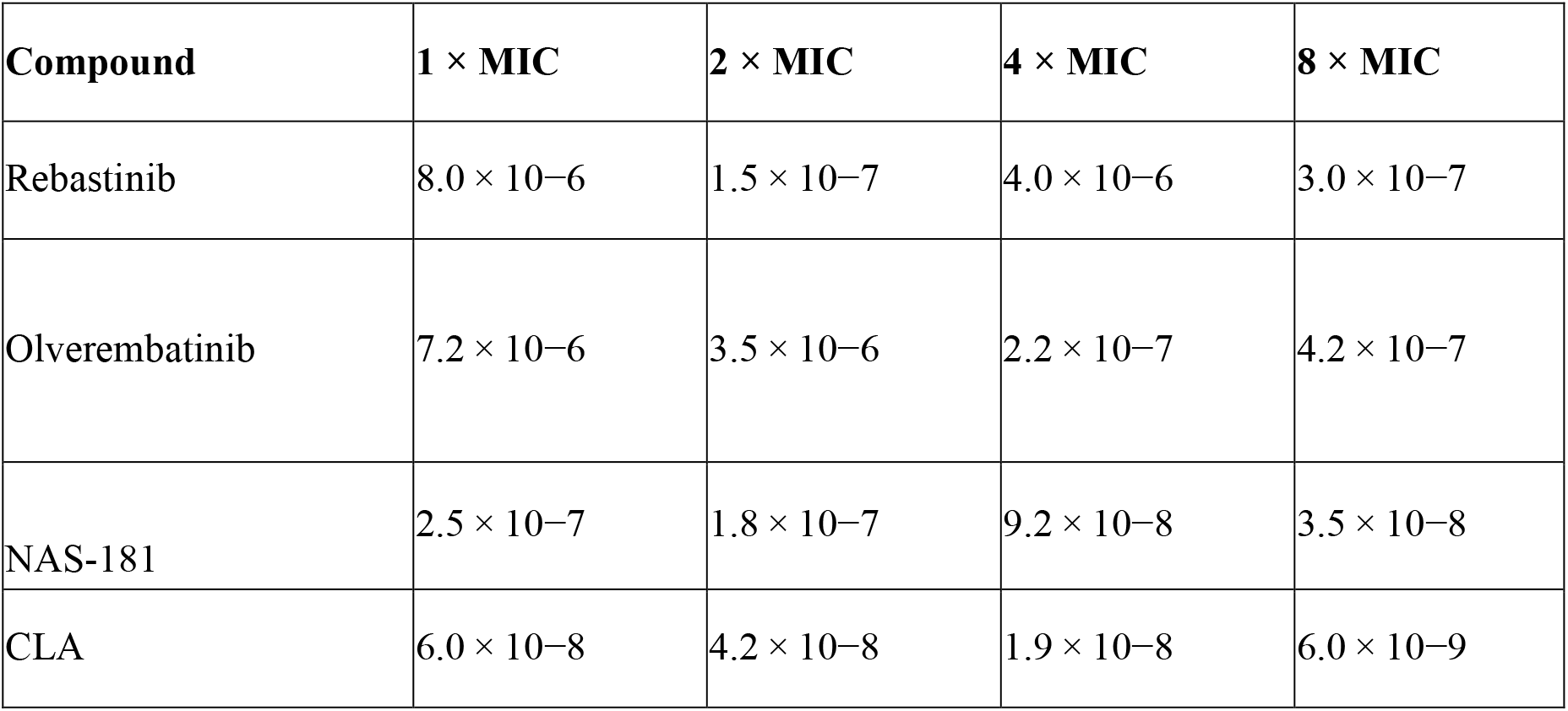
Resistance frequencies of *M. abscessus* ATCC 19977 to rebastinib, olverembatinib, NAS-181, and clarithromycin.

### 3.4. Checkerboard assay for compound interactions

To evaluate the potential of our hit compounds in combination with commonly used anti-*M. abscessus* agents, checkerboard assays were performed with rebastinib, olverembatinib, and NAS-181. First, the MIC_50_ values of all individual compounds were determined by REMA (Table 2). CLA showed favorable activity against *M. abscessus* ATCC 19977 (MIC_50_ = 0.5 µM), whereas the three hit compounds exhibited MIC_50_ values ranging from 9 to 40 µM.

Second, combinations were evaluated using 2-fold serial dilutions, and the interactions were interpreted by ΣFIC values (Table 3). The majority of combinations produced additive effects, with ΣFIC values between 2.0 and 3.4. Importantly, three combinations demonstrated clear synergy (ΣFIC ≤ 0.5): (i) rebastinib + moxifloxacin (ΣFIC = 0.40), (ii) olverembatinib + ceftibuten (ΣFIC = 0.35), and (iii) NAS-181 + clofazimine (ΣFIC = 0.35). These synergistic pairs required lower concentrations of both drugs to inhibit growth compared with the compounds used alone. As shown in Table 3, for example, one-half the MIC_50_ of rebastinib combined with one-half the MIC_50_ of moxifloxacin prevented resazurin turnover, consistent with synergistic growth inhibition.

**Table 3.**
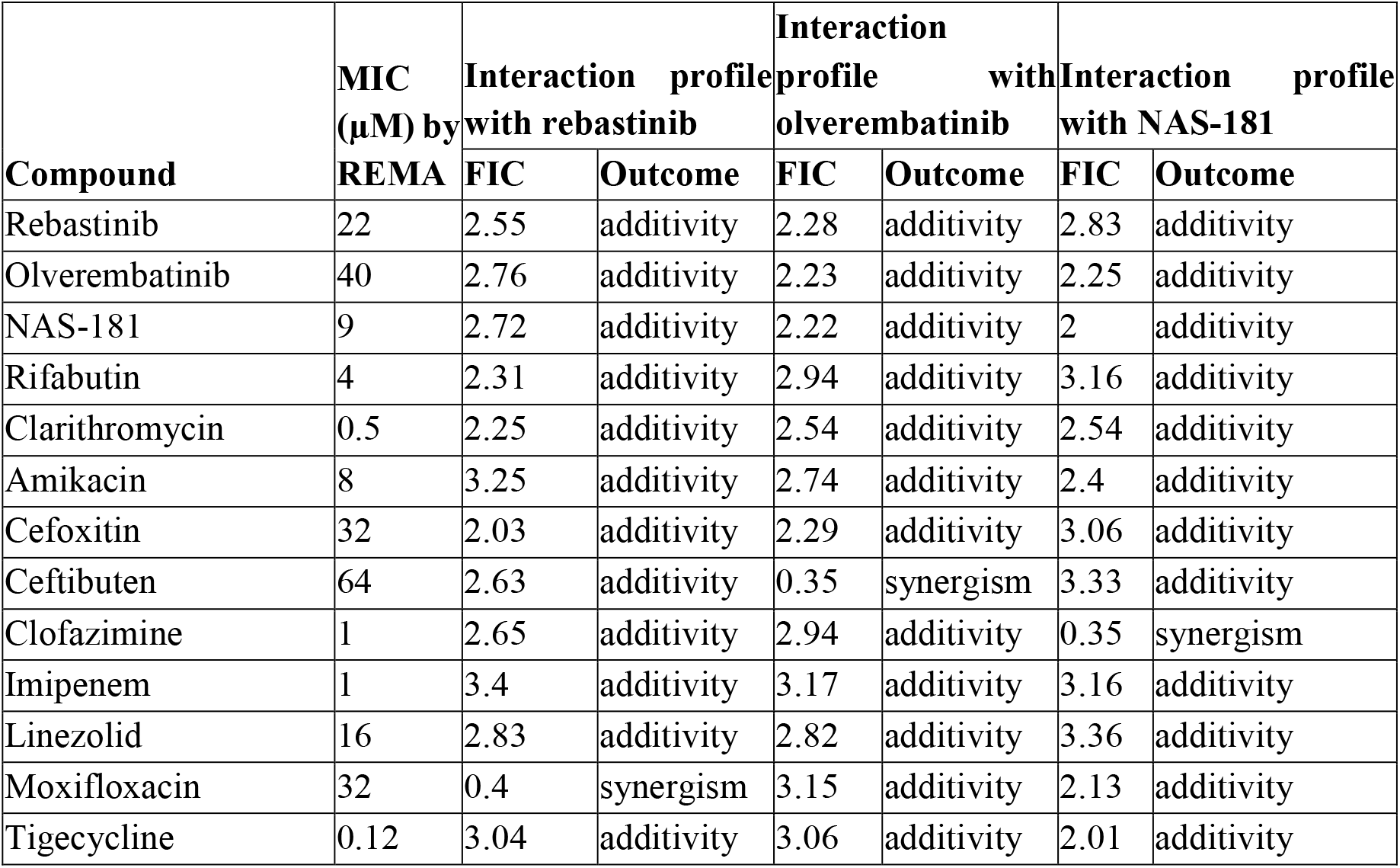
Drug–drug interaction profiles of rebastinib, olverembatinib, and NAS-181 with commonly used anti-*M. abscessus* agents.

Similarly, sub-MIC levels of olverembatinib with ceftibuten and NAS-181 with clofazimine also produced strong inhibitory effects. No antagonistic interactions were observed among any of the tested combinations.

Together, these findings indicate that while most interactions of the hit compounds with standard drugs were additive, specific combinations such as rebastinib–moxifloxacin, olverembatinib– ceftibuten, and NAS-181–clofazimine show synergistic potential and warrant further evaluation.

### 3.5. Target identification of rebastinib, olverembatinib, and NAS-181

To gain insight into the mechanisms of action of rebastinib, olverembatinib, and NAS-181, *M. abscessus* ATCC 19977 resistant mutants were isolated and analyzed. Resistant colonies emerged after exposure to each compound at 1× and 2× MIC, with resistance confirmed by REMA (MICs increased >8-fold relative to wild type). Whole-genome sequencing of representative resistant isolates revealed nonsynonymous single nucleotide polymorphisms (SNPs) in mshC (*MAB_2116*) and mshA (*MAB_4057c*), key genes in the mycothiol biosynthesis pathway. Specifically, one rebastinib-resistant clone carried a D87A substitution in MshC (*MAB_2116*), while an olverembatinib-resistant isolate harbored a W227G substitution in the same gene. In addition, a NAS-181-resistant mutant carried a Q51T substitution in MshA (*MAB_4057c*). These mutations were confirmed by Sanger sequencing in independent resistant isolates. Together, these findings strongly implicate the mycothiol biosynthesis pathway as a likely target of rebastinib, olverembatinib, and NAS-181, providing a mechanistic explanation for their antimycobacterial activity and supporting further investigation into mycothiol-related vulnerabilities in *M. abscessus*.

### 3.6. Activity of rebastinib, olverembatinib, and NAS-181 against intracellular replicating *M. abscessus*

First, the cytotoxicity of rebastinib (RES), olverembatinib (OLV), and NAS-181 against THP-1 cells was investigated over a 3-day exposure period. As shown in Figure S3, RES exerted significant cytotoxicity at concentrations above 30 μM, whereas OLV showed toxicity only at concentrations exceeding 60 μM. By contrast, NAS-181 displayed measurable cytotoxicity at a lower threshold of 14 μM, indicating higher host cell sensitivity compared with the other two compounds. Based on these results, all subsequent macrophage infection assays were performed using 20 μM rebastinib, 30 μM olverembatinib, and 7 μM NAS-181. Clarithromycin (CLA) at 0.1 μM was included as a positive control. Rebastinib, olverembatinib, and NAS-181 displayed potent intracellular activity against *M. abscessus* in THP-1 macrophages, with efficacy comparable to clarithromycin (CLA). As shown in Figure 1, DMSO-treated macrophages sustained high bacterial loads (∼10 log_10_ CFU/ml), whereas compound treatment led to significant reductions. Rebastinib and CLA each reduced intracellular survival by approximately 2 log_10_ CFU, while olverembatinib caused a moderate ∼1 log_10_ reduction. Notably, NAS-181 exhibited the strongest effect, decreasing intracellular bacterial counts by nearly 3 log_10_ CFU compared to the control. These findings demonstrate that all three compounds penetrate macrophages and effectively inhibit intracellular *M. abscessus*, with NAS-181 showing the most pronounced activity.

**Figure 1.**
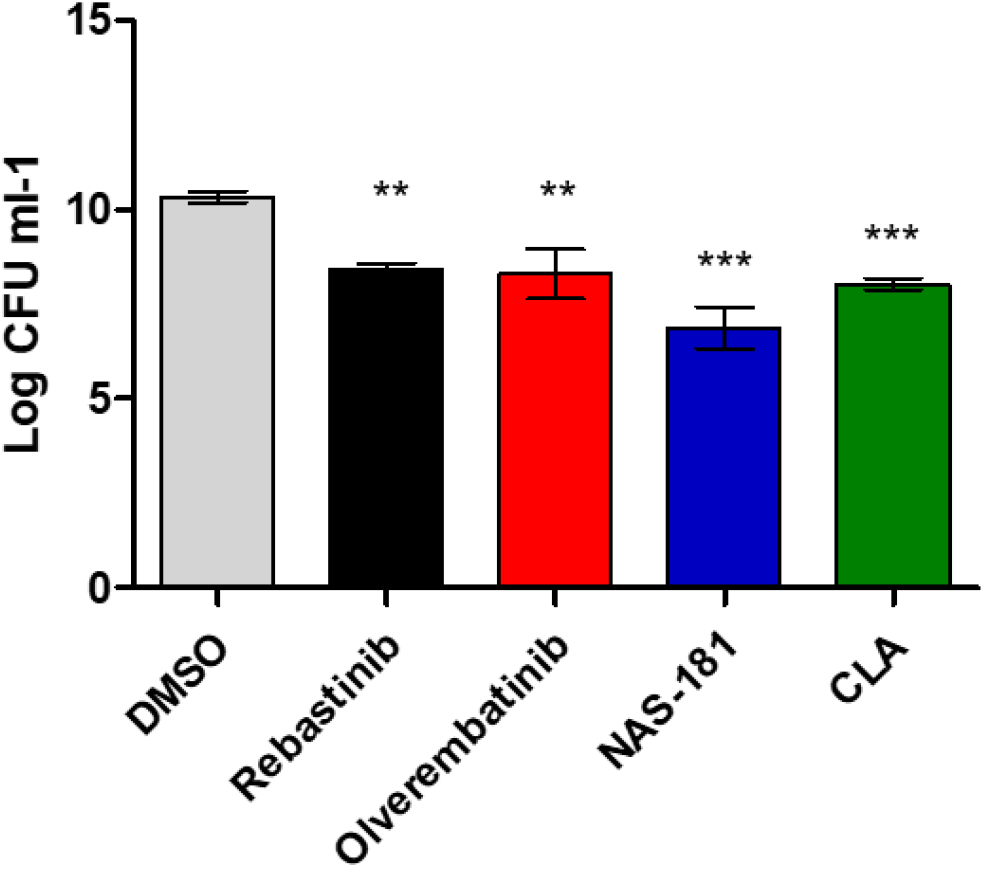
Intracellular activity of rebastinib, olverembatinib, NAS-181, and clarithromycin against *M. abscessus*. THP-1 macrophages were infected with *M. abscessus* at an MOI of 3 and treated with the indicated compounds. Intracellular bacterial survival was quantified by CFU enumeration after 72 h. The experiment was performed in triplicate, and the results are shown as the means ± standard deviation (SD). Rebastinib, NAS-181, and clarithromycin significantly reduced intracellular CFU compared to the DMSO control. ***, P < 0.001; **, P < 0.01.

### 3.7. Confirmation that hit compounds decrease mycothiol and GlcN-Ins levels

We next determined whether treatment with the hit compounds affected intracellular mycothiol (MSH) and its precursor GlcN-Ins (Table 4). The concentrations of MSH and GlcN-Ins were quantified in *M. abscessus* ATCC 19977 cells treated with rebastinib, olverembatinib, and NAS-181 at increasing concentrations (1×, 2×, 4×, and 8× MIC). Compared with the DMSO control, which yielded 25.2 nmol MSH and 1.8 nmol GlcN-Ins per 10^9^ cells, clarithromycin treatment showed no major effect on thiol levels (24.9 nmol MSH; 1.75 nmol GlcN-Ins). In contrast, all three hit compounds caused a progressive and concentration-dependent decline in both MSH and GlcN-Ins. Rebastinib reduced MSH from 20.2 nmol at 1× MIC to 12.4 nmol at 8× MIC, with GlcN-Ins levels decreasing from 1.35 to 0.82 nmol. A similar trend was observed with olverembatinib, which decreased MSH from 20.4 to 12.5 nmol and GlcN-Ins from 1.20 to 0.73 nmol across the tested range. NAS-181 treatment likewise resulted in a stepwise reduction, with MSH dropping from 18.6 to 11.4 nmol and GlcN-Ins from 1.15 to 0.70 nmol. These results demonstrate that rebastinib, olverembatinib, and NAS-181 significantly impair mycothiol biosynthesis in *M. abscessus* in a dose-dependent manner.

**Table 4.**
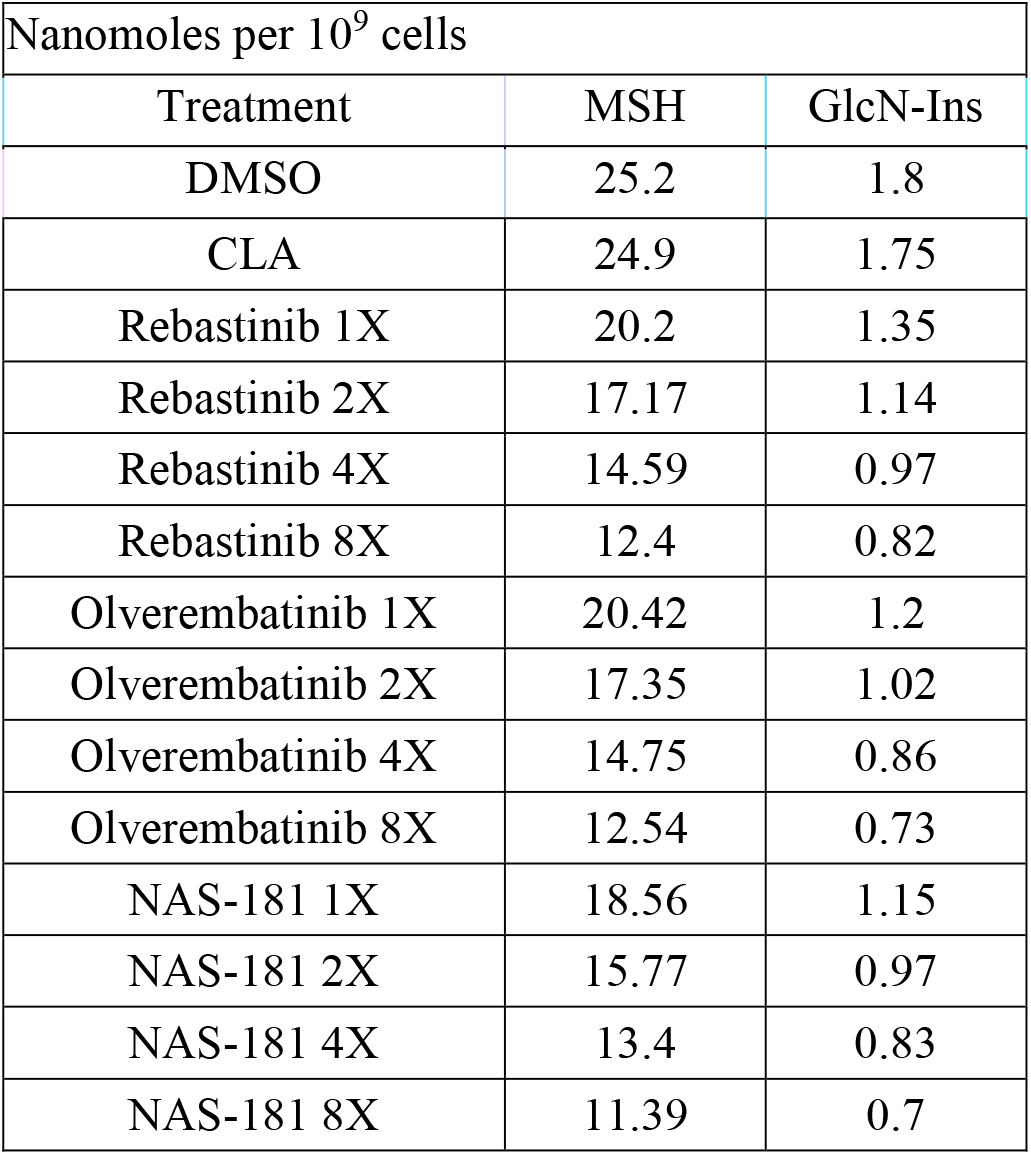
Effect of hit compounds on intracellular mycothiol (MSH) and GlcN-Ins levels.

| Nanomoles per 10<sup>9</sup> cells |  |  |
| --- | --- | --- |
| Treatment | MSH | GlcN-Ins |
| DMSO | 25.2 | 1.8 |
| CLA | 24.9 | 1.75 |
| Rebastinib 1X | 20.2 | 1.35 |
| Rebastinib 2X | 17.17 | 1.14 |
| Rebastinib 4X | 14.59 | 0.97 |
| Rebastinib 8X | 12.4 | 0.82 |
| Olverembatinib 1X | 20.42 | 1.2 |
| Olverembatinib 2X | 17.35 | 1.02 |
| Olverembatinib 4X | 14.75 | 0.86 |
| Olverembatinib 8X | 12.54 | 0.73 |
| NAS-181 1X | 18.56 | 1.15 |
| NAS-181 2X | 15.77 | 0.97 |
| NAS-181 4X | 13.4 | 0.83 |
| NAS-181 8X | 11.39 | 0.7 |

### 3.8. Efficacy of rebastinib, olverembatinib, and NAS-181 in a mouse pulmonary infection model

A neutropenic mouse pulmonary infection model was employed to evaluate the in vivo efficacy of our hit compounds against M. abscessus. Cyclophosphamide-pretreated mice were intranasally infected with ∼1 × 10^6^ CFUs of *M. abscessus* and subsequently treated with PBS, 5 mg/kg rebastinib, 5 mg/kg olverembatinib, 5 mg/kg NAS-181, or 50 mg/kg amikacin (AMK) as a positive control. Body weight monitoring revealed that all drug-treated groups maintained stable weight over the course of infection, whereas the PBS group exhibited marked weight loss beginning at day 3 and stabilizing at ∼80% of the initial weight (Fig. 2A).

**Figure 2.**
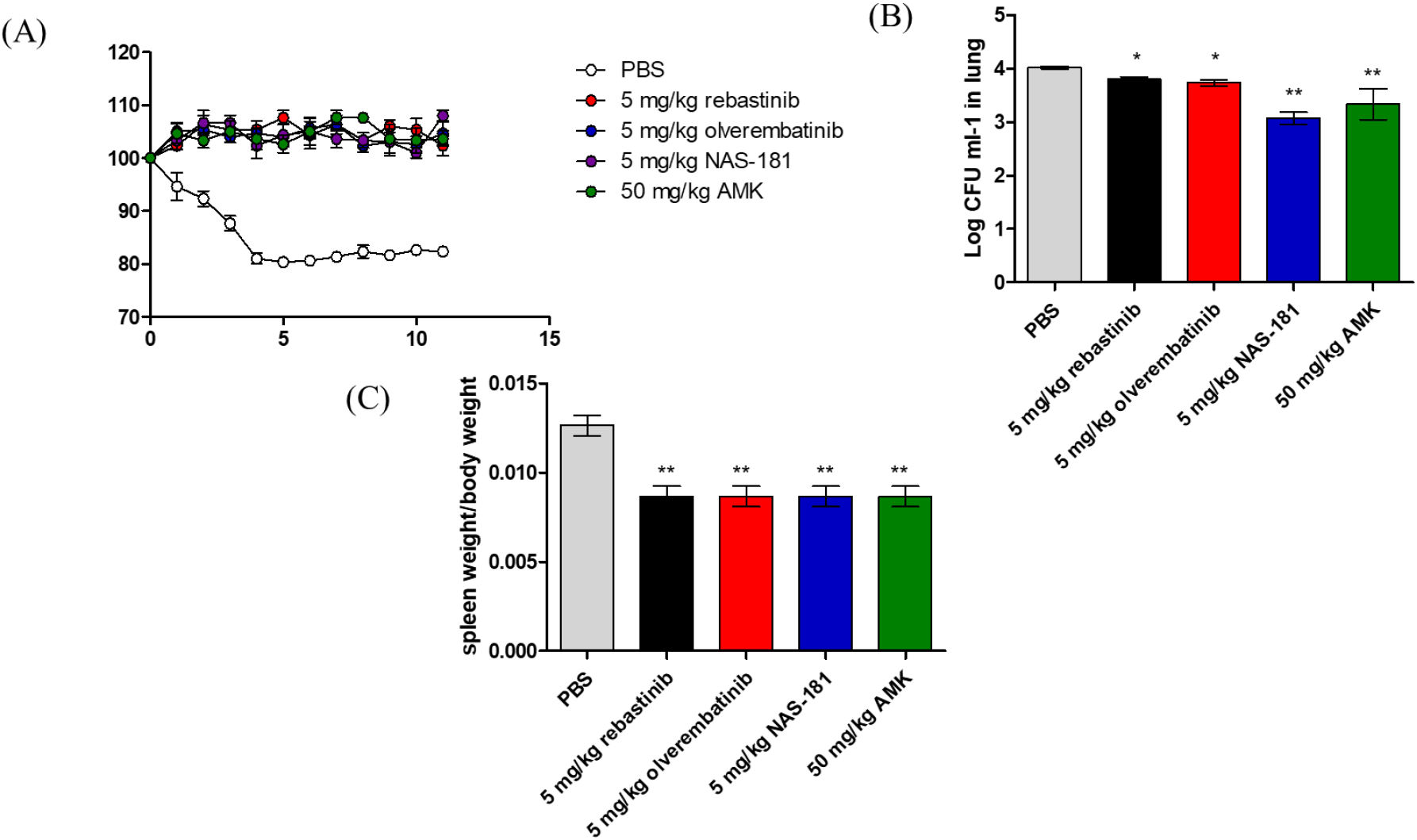
In vivo efficacy of rebastinib, olverembatinib, and NAS-181 in a mouse pulmonary infection model. (A) Body weight monitoring of neutropenic mice infected intranasally with *M. abscessus* (∼1 × 10^6^ CFUs) and treated daily with PBS (vehicle), 5 mg/kg rebastinib, 5 mg/kg olverembatinib, 5 mg/kg NAS-181, or 50 mg/kg amikacin (AMK). Data represent mean body weight change (%) ± SD (n = 5 per group). (B) Bacterial burden in lung homogenates at day 12 post-infection. Data are shown as mean log_10_ CFU/ml ± SD (n = 5). (C) Spleen enlargement expressed as the spleen weight/body weight ratio at day 12 post-infection. Data are shown as means ± SD (n = 5).

Bacterial burden analysis of lung homogenates showed that treatment with NAS-181 led to the most pronounced reduction in pulmonary CFUs, comparable to that achieved with amikacin (Fig. 2B). Rebastinib and olverembatinib also reduced bacterial burden relative to PBS, though to a lesser extent. Consistent with the CFU results, spleen enlargement was significantly attenuated in all treated groups compared with PBS, indicating alleviation of systemic infection (Fig. 2C).

Collectively, these data demonstrate that our hit compounds, particularly NAS-181, exhibit in vivo efficacy in reducing bacterial burden and infection-associated pathology in the *M. abscessus* mouse pulmonary model.

## 4. Discussion

In this study, we identified and characterized three kinase inhibitors, such as rebastinib, olverembatinib, and NAS-181 as novel candidates with antimycobacterial activity against *Mycobacterium abscessus* ^16–19^. All three compounds exhibited measurable inhibitory activity in vitro, with MICs ranging from 9 to 40 μM, and displayed predominantly additive interactions when combined with standard-of-care antibiotics. Notably, rebastinib demonstrated synergism with moxifloxacin, while olverembatinib and NAS-181 showed synergistic effects with ceftibuten and clofazimine, respectively, highlighting their potential as partners in combination regimens.

Genetic analysis of resistant mutants provided insight into the potential molecular targets. Single nucleotide polymorphisms were identified in genes associated with mycothiol biosynthesis, including *mshC* (D87A with rebastinib, W227G with olverembatinib) and *mshA* (Q51T with NAS-181). These mutations suggest that interference with the mycothiol pathway may underlie the antimycobacterial activity of these compounds. Consistent with this, biochemical quantification revealed that treatment with rebastinib, olverembatinib, or NAS-181 reduced intracellular mycothiol and its precursor GlcN-Ins in a dose-dependent manner, further supporting their activity against the MSH pathway.

Cytotoxicity testing confirmed that all three compounds were tolerated in mammalian cells, with toxic effects only observed at relatively high concentrations (30 μM for rebastinib, 60 μM for olverembatinib, and 14 μM for NAS-181). Importantly, intracellular assays using infected macrophages demonstrated significant reductions in bacterial load without compromising host cell viability, confirming that these compounds can penetrate host cells and act against intracellular *M. abscessus*.

Finally, in a neutropenic mouse pulmonary infection model, daily administration of rebastinib, olverembatinib, or NAS-181 (5 mg/kg) led to reduced bacterial burden in the lungs and ameliorated spleen enlargement compared with PBS-treated controls. Although the degree of bacterial clearance was less pronounced than with amikacin, all three compounds improved host weight stability and reduced disease-associated pathology, underscoring their in vivo efficacy. Together, these findings establish rebastinib, olverembatinib, and NAS-181 as promising leads against *M. abscessus*, acting at least in part through disruption of the mycothiol biosynthetic pathway. Their additive or synergistic interactions with existing antibiotics, coupled with intracellular and in vivo activity, highlight their potential for further optimization. Future studies should focus on structure–activity relationship analysis, detailed mechanistic studies on MSH pathway inhibition, and evaluation of combination regimens in more advanced animal models.

## CONTRIBUTOR CONTRIBUTIONS

D.L.S., E.M.D., and A.M. performed all experiments, including in vitro screening, MIC determinations, checkerboard assays, intracellular infection studies, mycothiol quantification, and the animal study.

A.P.S. and R.J.C. conceived and designed the study, coordinated the project, analyzed the data, and wrote the manuscript with contributions from all authors.

All authors reviewed and approved the final version of the manuscript.

## DATA AVAILABILITY

All data generated or analyzed during this study are included in this published article.

## ETHICS APPROVAL

The animal study was conducted in strict accordance with guidelines set forth by the Instituto de Biología y Medicina Experimental Animal Care and Use Committee. The study adhered to all ethical principles and regulatory standards for the care and use of laboratory animals.

